# High-quality rice RNA-seq-based co-expression network for predicting gene function and regulation

**DOI:** 10.1101/138040

**Authors:** Hua Yu, Bingke Jiao, Chengzhi Liang

## Abstract

Inferring the genome-scale gene co-expression network is important for understanding genetic architecture underlying the complex and various biological phenotypes. The recent availability of large-scale RNA-seq sequencing-data provides great potential for co-expression network inference. In this study, for the first time, we presented a novel heterogeneous ensemble pipeline integrating three frequently used inference methods, to build a high-quality RNA-seq-based Gene Co-expression Network (GCN) in rice, an important monocot species. The quality of the network obtained by our proposed method was first evaluated and verified with the curated positive and negative gene functional link datasets, which obviously outperformed each single method. Secondly, the powerful capability of this network for associating unknown genes with biological functions and agronomic traits was showed by enrichment analysis and case studies. Particularly, we demonstrated the potential applications of our proposed method to predict the biological roles of long non-coding RNA (lncRNA) and circular RNA (circRNA) genes. Our results provided a valuable data source for selecting candidate genes to further experimental validation during rice genetics research and breeding. To enhance identification of novel genes regulating important biological processes and agronomic traits in rice and other crop species, we released the source code of constructing high-quality RNA-seq-based GCN and rice RNA-seq-based GCN, which can be freely downloaded online at https://github.com/czllab/NetMiner.

## Introduction

The complex cellular network formed by the interacting macromolecules underlie an organism’s phenotypes (Kitano, 2002a, 2002b; Vidal et al., 2011). Reconstructing a complete map of the cellular network is crucial for understanding an organism’s genetic architecture underlying phenotypes. In animals, multiple types of networks have been built based on multi-level ‘-omics’ datasets from genome, transcriptome, proteome,epigenome, metabolome and other subcellular systems (Mitra et al., 2013). In plants, most of the current available ‘-omics’ dataset comes from the transcriptome analysis, with relatively few studies generating other types of ‘-omics’ datasets (Ma et al., 2013). The rapid accumulation of large-scale open access plant transcriptome data provides the great potential for identifying the molecular networks underlying diverse functions. Co-expression meta-analysis is a powerful method for reconstructing gene co-expression network using transcriptome data. This method combines expression profiles from all available experimental conditions, aims to predict the statistically significant functional associations between genes. The extensibility and easiness to apply make it a powerful tool for inferring the biological roles of uncharacterized genes (Bergmann et al., 2003; Gerstein et al., 2014; Ma et al.,2013; Mutwil et al., 2011; Stuart et al., 2003).

For co-expression meta-analysis, many algorithms have been proposed to construct the gene networks. However, it has been shown that the outcome of network inference varies between tools, and the single network inference approach has inherent biases and is unable to perform optimally across all experimental datasets (De Smet and Marchal, 2010; Marbach et al., 2012). In addition, how to clean-up the links occurring by accident in a gene co-expression network and select biologically significant associations is also a critical procedure for modeling the authentic gene relations (Alipanahi and Frey, 2013; Usadel et al., 2009). Moreover, the current computational methods are mainly designed for analyzing microarray dataset. Indeed, microarrays are intrinsically limited for measuring a relative small dynamic range of gene expression and only representing a subset of genomic contents (Abdullah Sayani et al., 2006; Mutwil et al., 2011). Compared with microarrays, RNA sequencing (RNA-seq) emerges as a new approach to transcriptome profiling, which provides broader dynamic range of measurements allowing genome-wide detection of novel, rare and low-abundance transcripts. However, the majority of co-expression meta-analyses have been neglected the rapid growing availability of next-generation RNA-seq data (especially in plants). Its potential capacity in co-expression network inference has not been well studied.

In this study, we designed a novel ensemble pipeline for inferring high-quality Gene Co-expression Network (GCN) using RNA-seq data by integrating the predictions of three different network inference algorithms. Since the multiple types of networks in the model plant, *Arabidopsis,* has been constructed and widely analyzed, we directly applied this pipeline to the important crop species, rice, to enhance its efficiency of molecular breeding. We compiled a standard physical and non-physical set of positive and negative functional link datasets between genes derived from 4 known biological networks and evaluated the quality of our network. In the case study, bottom-up subnetwork analysis revealed that the usefulness of reconstructed RNA-seq-based gene co-expression network for realistic biological problems. Particularly, we showed that the potential application of our method for predicting the biological roles of the uncharacterized genome elements including long non-coding RNA (lncRNA) and circular RNA (circRNA) genes. Our study revealed the massive genetic regulatory relationships associating with cellular activities and agronomic traits, which provide a valuable data source for selecting candidate genes to accelerate rice genetics research.

## Results

### Network construction and evaluation

To evaluate the quality and reliability of publicly available RNA-seq dataset, we analyzed 348 RNA-seq transcriptomes of the important monocot crop species rice after removing the unreliable genes and samples (for details, see Dataset 2, Materials and methods section). After quality filtering and trimming, a total of 12,458,505,209 reads were remained in the samples, 75.2% of which were mapped to the MSU7.0 reference genome and 71.4% were mapped uniquely (see Dataset 2). Of the genes (MSU7.0 reference set) covered with RNA-Seq reads, 98.4% have coverage of > 50% of the gene length (see Supplementary Information, Fig.S1A). Despite of the large difference in the number of mapped reads between samples, the percentage of expressed genes is similar in most of them, ranging from 32% (10th percentile) to 66% (90th percentile), and as the number of mapped reads increases, the ratio of the number of expressed genes is rapidly increased to saturation (see Supplementary Information, Fig.S1B). We tested several normalization methods to compute the expression abundance and expression correlations between genes and samples, the tissue-specific expression pattern and enrichment results of rice genes showed that these RNA-seq data are highly reliable (see Supplementary Text, Fig.S2-Fig.S6, Table S1 and Dataset 3 for details).

We comprehensively analyzed whether the co-expression between genes is associated with their biological roles, and demonstrated that functionally related genes are often to be co-expressed in our RNA-seq dataset (see Supplementary Text, Fig.S7-Fig.S8, Dataset 4 for details). Based on this, we designed a new ensemble pipeline to build RNA-seq-based gene co-expression network by integrating the predictions of three state-of-the-art network inference methods, including Weighted Gene Correlation Network Analysis (WGCNA) (Langfelder and Horvath, 2008), Graphical Gaussian Model (GGM) (Schäfer et al., 2001) and Bagging the Conservative Causal Core of Network (BC3NET) (de Matos Simoes and Emmert-Streib, 2012), based upon an un-weighted voting system and rescoring the co-expression links (see Materials and Methods for details). We here did select these three inference methods but not the other existing approaches is because of either their high computational complexity or the inconsistent data source (Feizi et al., 2013; Friedman et al., 2008; Huynh-Thu et al., 2010; Qin et al., 2014). We constructed the co-expression network of rice which included 16770 genes with 146,419 links. This network shows the small-world characteristic with an average path length between any two nodes is equal to 6.28. The distribution of connection degrees obeys the truncated power-law where most nodes have a few co-expression partners with only a small ratio of hub nodes associating with a large number of partners (see Supplementary Information, Fig.S9A). The negative correlation between degrees and clustering coefficients of genes reveal hierarchical and modular characteristics of network and the possible synergistic regulation of gene expression (Supplementary Information, Fig.S9B) (Bergmann et al., 2003).

We evaluated the performance of the ensemble inference pipeline in rice. Since there are no gold standard reference co-expression networks available in rice, we compiled as replacement a standard set of positive links (9390203 interactions), by capturing gene pairs that were contained in the same Gene Ontology (GO) categories, the same pathways, interact with each other in the protein-protein interaction network or linked in the probabilistic functional gene network (RiceNet), and a standard set of negative links (272997 interactions) based on the functional dissimilarities between genes (for details, see Materials and methods section). We used fold enrichment to measure the relationship of two data sets (our network and standard positive functional links / our network and standard negative functional links): the larger the proportion of the number of shared elements divided by that expected by random chance, the closer they are (see Materials and methods for details). We found that the co-expression relationships connecting highly or frequently expressed gene pairs were positively associated with the positive standard links and were negatively associated with the negative standard links (see Supplementary Information, Fig. S10). Meanwhile, we also observed that the expression sample number of co-expression link (defined as the total number samples which simply plus the number of gene A expressed samples and the number of gene B expressed samples) is a more reliable factor than its expression level (defined as the expression abundance summation of gene A and gene B) to affect the fold enrichment of the standard links (see Supplementary Information, Fig. S10). These outcomes indicated that the positive standard links had reliably captured the co-expression links between genes. Using the standard datasets, we found that the network structure obtained by our ensemble inference method was consistently better than the networks built by the individual method with higher enrichment for positive links and lower enrichment for negative links (Fig.1). These results suggested that the committee of different methods can reduce the bias occurring in a single inference method and provide more reliable predictions with higher sensitivity and specificity. We observed that the folds of enrichment are not obviously improved or are slightly decreased by the integrated networks from 6 data set (Fig.1A, the GGM method, line highlighted in yellow) than that of each single data set, indicating that integrating the networks built using different data normalization methods might have no obvious effects on the structure of inferred network (Fig.1). Co-expression is actually one of the inputs used to build the probabilistic functional gene network (RiceNet), which were included in the standard positive links. To examine whether this has effect on our evaluation results, we carried out the fold enrichment analysis after removing the links contained in RiceNet from the standard positive links. We found that integrating the functional links of RiceNet into the standard positive links has no effect on the results of comparing the quality of our network with the other networks obtained by the single algorithm (see Supplementary Information, Fig. S11). Based on the novel RNA-seq dataset, we also examined whether a large fraction of potential interactions was recovered by our collected RNA-seq dataset, and found that the most general transcriptional links were already established reliably with these 348 rice RNA-seq samples (see Supplementary Text for details).

**Fig 1.**
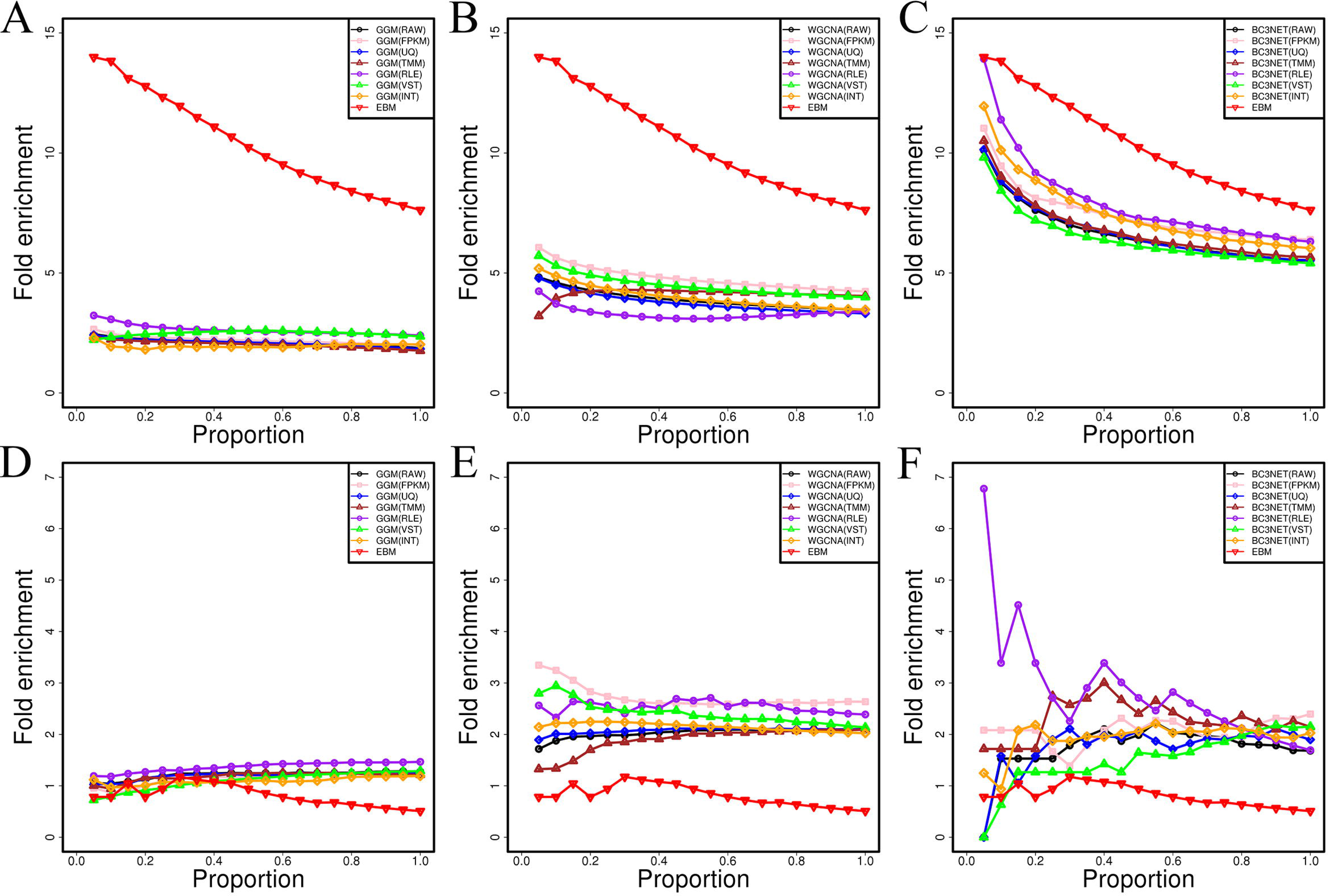
Enrichment folds of different algorithms for co-expression network inference. A) Comparing to GGM with positive links. B) Comparing to WGCNA with positive links. C) Comparing with BC3NET with positive links. D) Comparing with GGM with negative links. E) Comparing with WGCNA with negative links. F) Comparing with BC3NET with negative links. In the legends, the RAW, FPKM, UQ, TMM, RLE and VST represent the networks obtained by the single RNA-seq dataset; INT indicates intra-method consensus networks established by integrating the predictions ofdifferent RNA-seq datasets, EBM denotes high-quality gene co-expression network obtained by integrating all intra-method consensus networks

### Prediction of gene functions through co-expression subnetworks

We observed that our reconstructed RNA-seq-based gene co-expression network is always positive predictor of functional associations for the protein-protein interaction network and probabilistic functional gene network, GO network and pathway network (see Supplementary Text, Fig. S12). Meanwhile, we also observed that many genes under the same GO functional category are significantly more connected to each other than expected by chance (see Supplementary Text, Dataset 5). Therefore, we adopted GO enrichment analysis of a gene’s co-expression neighborhood as a tool to predict its biological functions (Vandepoele et al., 2009). For each gene belonging to a given GO category, we asked whether the GO enrichment in its co-expression neighborhood could infer its correct function: an inference is called true positive if and only if the predicted GO term is more specific than its known GO terms or is equal to the known GO terms. In the enrichment significance level of corrected *p*-value smaller than 0.05, we found that 15.50% (Sensitivity) annotated functions were correctly inferred based on 10545 annotated genes in rice network. If we used only the gene annotations on the second and third layers of the directed GO graph for inference, the Sensitivity was increased to 21.66%. We found that the 21.27% (Precision) of all inferred functions are true positives and this number is improved to 25.38% when we only adopted the second and third layers of directed GO graph. These results might be suggesting that the incompleteness or errors in the GO annotations of rice genes.

The relatively low Sensitivity and Precision of our network in function inference might be due to the simple scoring metrics. We here further analyzed the predictive performance of our network based on the Critical Assessment of protein Function Annotation (CAFA) metrics (Tzafrir et al., 2003) (see Materials and Methods). To eliminate the effects of the incompleteness and errors of GO annotations, we removed the genes with I) the number of known annotations smaller than 3; II) the number of predicted annotations smaller than 3 and III) the variation coefficient of the number of known annotations and the number of predicted annotations larger than 0.5. To order to produce the Receiver Operating Characteristics (ROC) and Precision-Recall (PR) curves, we calculated the sensitivities, 1-specificities and precisions under different thresholds (-log(corrected *q*-value)). For the purpose of correcting different depths of GO predictions, we also calculated the weight value of each GO term and obtained the weighted ROC and PR curves. The weighted ROC and PR curves obtain the larger AUC score (70.01%) and maximum F-measure (F-max = 0.54) than the not weighted ones(AUC = 68.23%, F-max = 0.53) (see Fig.2), indicating that our gene network can effectively predict the difficult or less frequent GO terms (see Fig.2). In addition, we further compared the predictive performances of our network with RiceNet using the same evaluation criteria as employed in our study. We observed that our co-expression network is comparable or better than the RiceNet in terms of the ROC and PR curves (Fig.2). Moreover, we also found that the semantic similarities between the known GO terms and our predicted GO terms are obviously higher than the random ones (*p*-value = 5.24E-10, paired t-test). These results indicated that our RNA-seq-based gene network can be applied for inferring the potential functions of unknown genes.

**Fig 2.**
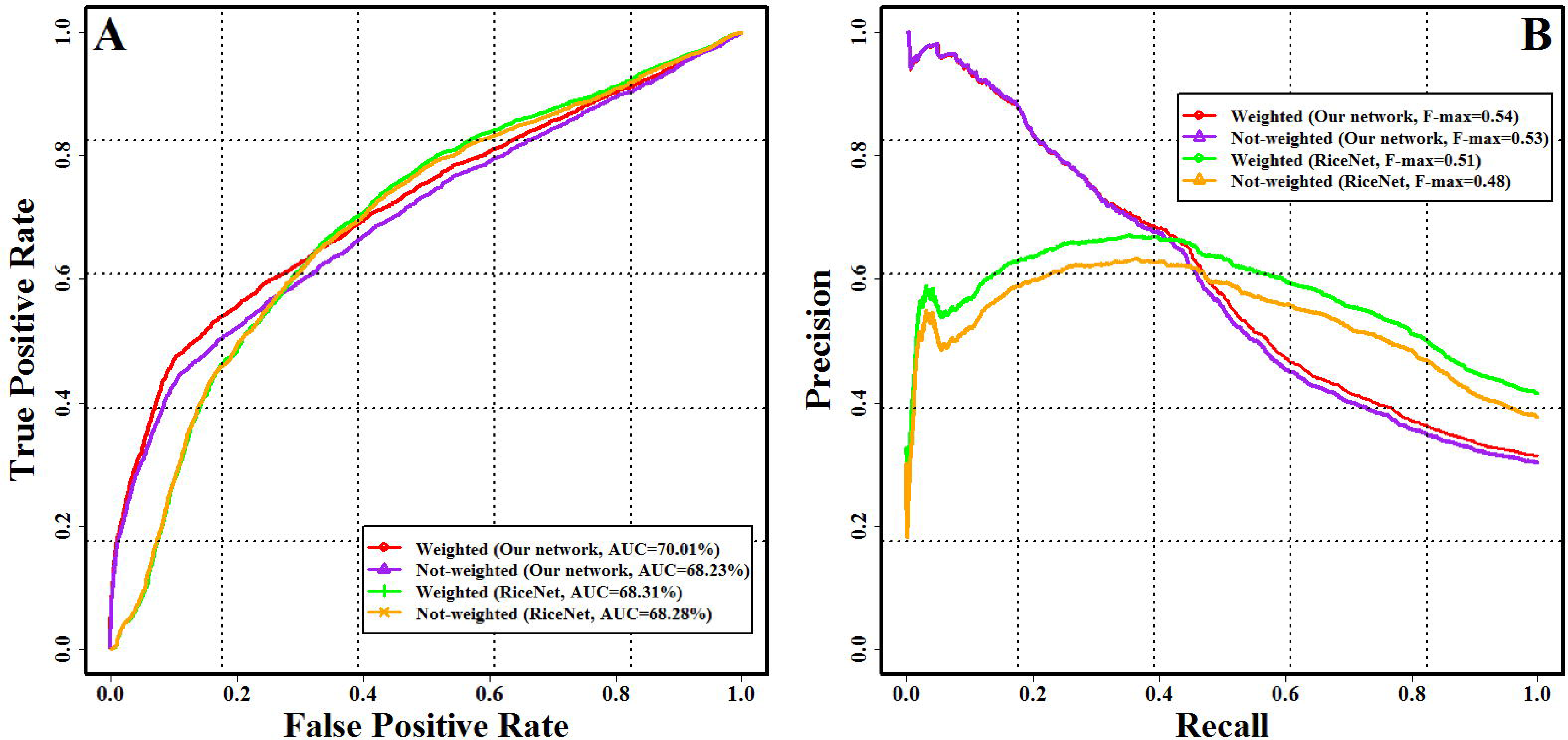
Performance evaluation of our network for predicting gene function. A) Receiver Operating Characteristics (ROC) curve. B) Precision-Recall (PR) curve. In the legends, Not-weighted indicates the evaluation parameters were calculated by the standard method of CAFA project; Weighted indicates the evaluation parameters were calculated by the weighted method of CAFA project

In addition to the neighboring gene analysis above, we used two examples below to demonstrate the stricter and intuitive method of RNA-seq-based gene co-expression network analysis for inferring the gene functions. In flowering plants, floral organ development is a very important biological process. We therefore first selected a priori guide gene *OsMADS16* involving in flower development to obtain a co-expression subnetwork consisting of 37 closely connected neighbors within two-layer links from the guide genes (see Fig.3A and Dataset 6). We found that 15 genes were involved in flower development process, with ~ 203-fold enrichment. For example, 11 members of MADS-box family, which were verified involving in the determination of floral organ identity and development, are effectively captured in this subnetwork. Moreover, this subnetwork includes the well-known genes *DL, Wda1* and *DPW,* which have been experimentally validated to control the floral organ identity, anther and pollen development (Jung et al., 2006; Nagasawa et al., 2003; Shi et al., 2011). Interestingly, we did not find that two YABBY domain containing genes *OsYABBY1* and *OsYABBY6* are annotated involving in floral organ development in rice, but their *Arabidopsis* homologs of *YABBY2* and *YABBY1* were associated with the inflorescence meristem growth and regulation of floral organ development (Siegfried et al., 1999). The connections between the unannotated genes (gray nodes) and known genes within a subnetwork provide clues for their associations with specific biological processes. For example, *LOC_Os07g09020* involves in the reproduction and embryo development, whose links with *OsMADS3, OsMADS4* and *DL* enable further targeted experimental validations.

Second, we used another guide gene *OsCESA4* involving in cell wall metabolism to build a subnetwork (Fig.3B and Dataset 6). The resulting subnetwork was made up of 139 genes with ~96-fold enrichment, including 4 homologs of *OsCESA4: OsCESA1, OsCESA3, OsCESA7* and *OsCESA9,* and 14 other genes associated with the cell wall metabolism. In addition, this subnetwork also captures 28 genes (pink nodes) whose *Arabidopsis thaliana* homologs were involved in cell wall metabolism. For example, *LOC_Os01g06580,* encoding a fasciclin domain containing protein, is a homologous gene to *AT5G03170* which is involved in secondary cell wall biogenesis. Two genes of *LOC_Os01g62490* and *LOC_Os03g16610* are laccase precursor proteins are both homologs to *LAC17* involved in cell wall biogenesis. *AT1G09540,* an *Arabidopsis* homolog of two rice MYB family transcription factors of *LOC_Os05g04820* and *LOC_Os01g18240,* are participating in cell wall macromolecule metabolism and xylem development. We also noted that 14 genes labeled with blue nodes, involving in carbohydrate metabolism, associating with microtubule or resembling to known cell wall metabolism genes in function domain, are recovered in this gene subnetwork. All these genes are the potential candidates for the further functional investigation. Especially, the known cell cycle genes *LOC_Os04g28620* and *LOC_Os04g53760* are also captured in this subnetwork, confirming that cell wall metabolism and cell cycle are two closely associated processes.

**Fig 3.**
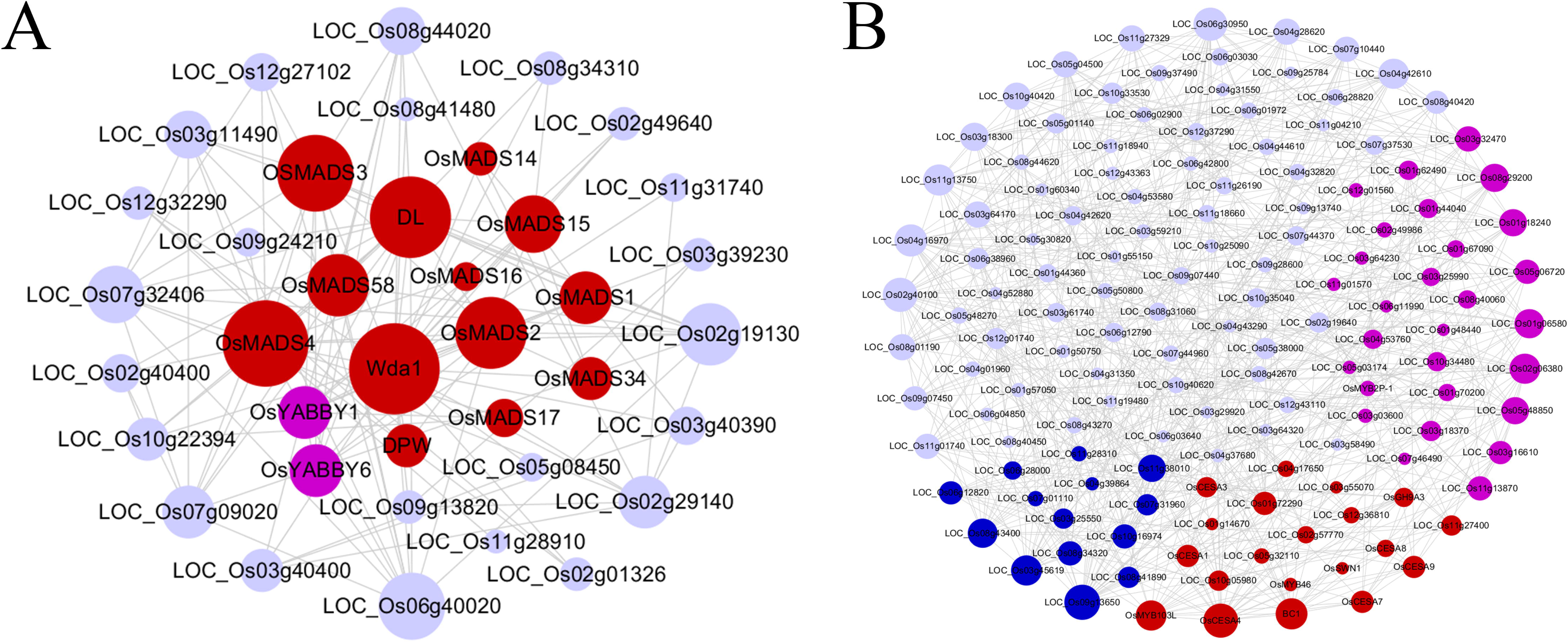
Subnetworks derived from the gene-guide approach. The subnetworks include all other nodes within twolayer connections from guide genes. A) *OsMADS16* involved in flower development; B) *OsCESA4* involved in cell wall biosynthesis. Within each subnetwork, red nodes represent the experimentallyverified genes related to corresponding biological functions. Pink nodes indicate the genes whose *Arabidopsis* homologs are experimentally verified relating to the corresponding biological processes. Blue nodes represent potential function-related genes, and the gray nodes denote that the genes with unknown functions or annotated with irrelevant functions. The size of node is proportional to the number of connected genes

### Construction of regulatory subnetworks for gene functional analysis

We explored the potential value of motif-guided analysis (Ma et al., 2013) in building regulatory network and finding functionally related genes using two examples. Cell cycle is a highly conserved biological process in higher eukaryotes. From G1 phase to S phase of the cell cycle is controlled by the E2F transcription factors, which bind to a conserved DNA motif WTTSSCSS (with “W” standing for “A” or “T” and “S” standing for “C” or “G”) (Vandepoele et al., 2005). We used this motif to retrieve 1093 genes from the rice network. Out of the 180 cell cycle genes annotated in rice (totally 55986 genes), 33 cell cycle genes were included in these 1093 genes, resulting in 9.4-fold enrichment. We used the cell cycle genes and the genes that were directly linked to them to form a regulatory network (totally 104 genes, Fig.4A and Dataset 6). We observed that a large number of genes (red nodes in Fig.4A) encode proteins participating in regulation of cell cycle, DNA replication, chromatin dynamics and DNA repair. The currently known cell cycle genes include three cyclin genes, one E2F transcription factor, 9 DNA replication origin factors, two checkpoint regulators, 13 DNA replication or repair proteins and 10 other genes with unknown biochemical functions but were annotated playing important roles during cell cycle. In addition, this subnetwork also includes 18 genes whose *Arabidopsis* homologs participate in regulation of cell cycle, DNA replication, DNA repair and chromatin dynamics. Also recovered are four genes including *LOc_Os01g64900, LOc_Os03g49200, LOC_Os07g18560* and *LOC_Os09g36900* whose *Arabidopsis* homologs have not annotated biochemical function but were involved in cell cycle. Although some genes are not annotated with direct participation of cell cycle, their molecular structure and function domain indicated their potential roles in it, such as the ribonuclease H2 subunit B *(LOC_Os04g40050),* ATP-dependent RNA helicase *(LOC_Os11g4491ű),* ribonuclease H2 subunit B *(LOC_Os04g40050)* and the BRCA1 C Terminus domain containing protein *(LOC_Os08g31930).* All these genes are the potential candidate cell cycle genes for further investigation.

WRKY transcription factors play important roles in regulation of plant stress response by binding the W-box sequence TTGACY (with “Y” standing for “C” or “T”) (Chen et al., 2012;Rushton et al., 2010). Similarly, we extracted a total of 1329 genes associating with W-box, from which a subset of 88 known stress response genes out of 996 genes relating to stress response in rice were found, achieving the fold enrichment of 3.72. We also constructed a regulatory network using the 88 genes and the genes with W-box that were directly linked to them (totally 389 genes, Fig.4B and Dataset 6). This subnetwork includes 172 genes that are regulated by different types of environmental stresses (red node). Among them, 138 rice genes and 34 homologs in *Arabidopsis* are annotated in the reference genomes relating to abiotic and biotic stresses. The majority of *Arabidopsis* homologs of these genes are experimentally confirmedinvolving in the biological regulation of phosphate starvation, water deprivation, nitrate, hypoxia, salt, cold, heat, chitin, sugar and oxidative stresses. Particularly, 53 of 172 abiotic stress response genes whose *Arabidopsis* homologs are reacted to the ethylene (ETH), abscisic acid (ABA), salicylic acid (SA) or jasmonic acid (JA), which is in accordance with the fact that WRKYs play roles in the plant abiotic stress by invoking the ETH-, ABA-, SA- or JA-mediated signaling pathways (Chen et al., 2012). Moreover, 36 genes play important roles in regulating plant immune responses to pathogens including WRKYs, NB-ARC domain containing resistance proteins, NBS-LRR domain containing resistance proteins, kinase proteins and other verified defense members of the plant innate immune system were also contained in this network (see Dataset 6). This is completely supported by the transcriptional reprogramming network model of the WRKY-mediated plant immune responses (Eulgem and Somssich, 2007). In addition, this gene subnetwork also included 8 genes whose *Arabidopsis* homologs are associated with the seed development, dormancy and germination. In agreement with the fact that the SA and ABA antagonizes gibberellin (GA)-promoted seed germination; 6 of these genes participate in the SA-and ABA-mediated signaling pathways (Xie et al., 2007). Interestingly, three genes of *LOC_Os03g12290, LOC_Os01g24550* and *LOC_Os01g 64470*involving in leaf senescence are also placed in this network, with *LOC_Os 01g 64470* involving in the SA- and JA-mediated signaling pathway, which is supported by the fact that the WRKYs function in leaf senescence by modulating the JA and SA equilibrium (Miao and Zentgraf, 2007). This subnetwork successfully captured the W-box related genes that can facilitate further studies the functions of uncharacterized genes and help us to understand the regulatory mechanisms of plant responding to various stresses.

**Fig 4.**
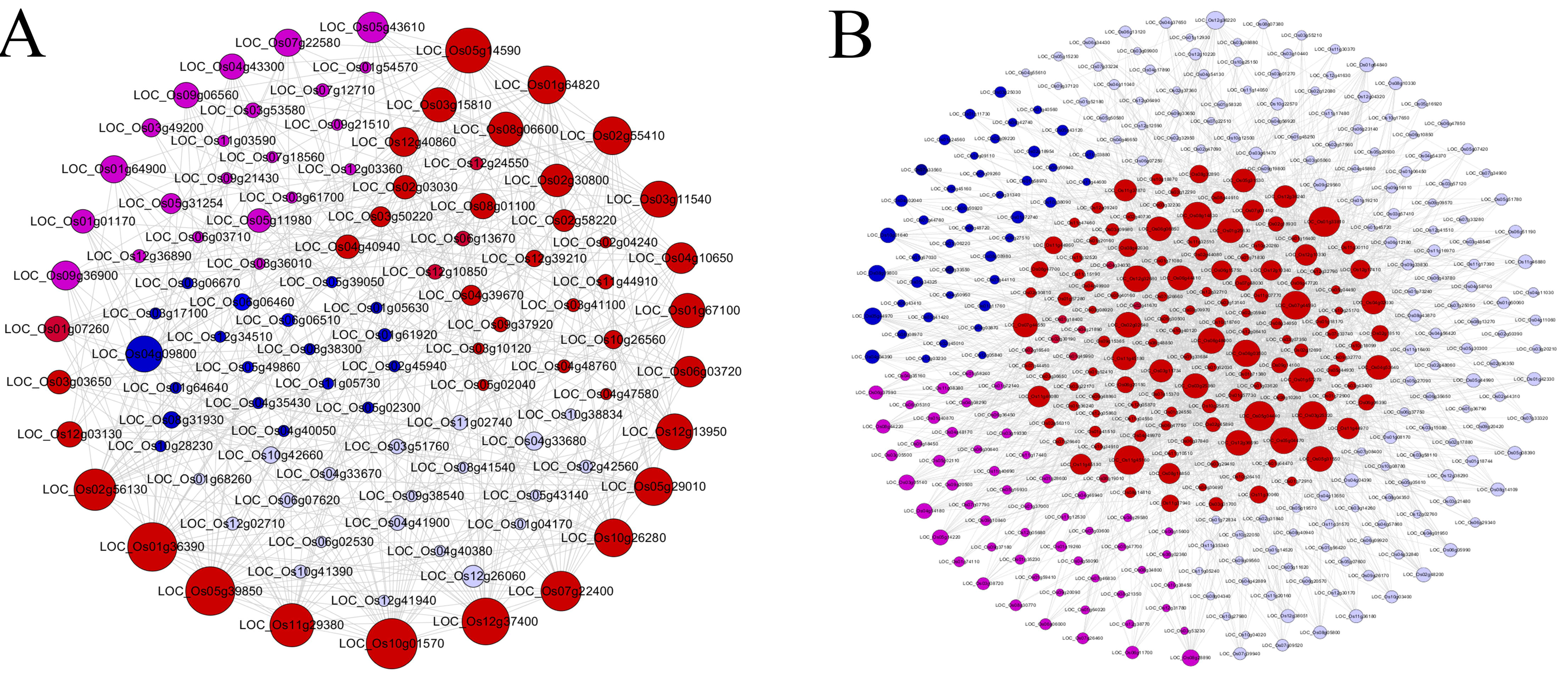
Subnetworks derived from the known cis-regulatory motif-guide approach. A) WTTSSCSS combined with the E2F transcription factors involved in cell cycle. B) TTGACY combined with the WRKY transcription factors involved in stress response. Within each subnetwork, red nodes represent the experimentally verified genes related tocorresponding biological functions. Pink nodes indicate the genes whose *Arabidopsis thaliana* homologs are experimentally verified to associate with the corresponding biological functions. Blue nodes denote potential function-related genes. Gray nodes indicate that the genes with unknown functions or annotated with irrelevant functions. The size of node is proportional to the number of connected genes

In addition, we also used two miRNAs of osa-miR156 and osa-miR396 to capture the functionally related genes based on microRNA target enrichment analysis, which is performed similar with motif enrichment analysis (Ma et al., 2013). We observed that a large number of genes involving in cell division and organ development were captured in this gene subnetwork, for example, two TCP transcription factors of *LOC_Os01g55100* and *LOC_Os11g07460* (see Fig.S13 and Dataset 6). Meanwhile, we also found that many genes relating to stress tolerance were placed in the subnetwork of osa-miR156, for instance, a WRKY transcription factors *LOC_Os 10g18099* (see Fig.S13 and Dataset 6). These obtained results well confirm the biological roles of these two miRNAs (Rodriguez et al., 2010; Stief et al., 2014; Wu et al., 2009). Taken together, all these outcomes indicated that the rice RNA-seq-based gene co-expression network could be converted to highly reliable regulatory network for further studying gene regulations.

### Co-expression analysis of genes controlling the important agronomic traits

For the perspective of system biology, the phenotype of an organism was controlled by functionally linked genes involving in the related biological processes. Given the co-expressed genes tend to have the related biochemical functions; we next want to use the co-expression relationships between genes to assign the agronomic traits for unknown genes. This is especially important for identifying the candidate genes in Quantitative Trait Loci (QTL) mapping, Genome-Wide Association Study (GWAS) or in reverse genetic studies. We collected 1031 known rice genes with the well-studied functions through wet lab experiments. For these genes, we found that 934 genes were expressed in our collected RNA-seq datasets and 623 genes were in network with 12125 connections. To examine the potential capacity of our RNA-seq-based gene co-expression network for associating genes with the agronomic traits, we analyzed the density of co-expression links between genes of within and between agronomic traits. We found that 262 co-expression links out of 88041 all possible links within the common agronomic traits and that 252 co-expression links out of 982302 all possible links between the different agronomic traits were captured in network, with ~11-fold enrichment of links within the agronomic traits. In details, we found that several agronomic traits whose genes were tightly clustered together relative to the average link density of whole co-expression network (Supplementary Text, Table S2). For example, an agronomic trait, source activity, measuring the capacity of making photosynthetic products; whose genes was highly aggregated in network with the enrichment fold of 47.81 and the corrected *p*-value of 3.96E-117. Besides, genes associating with culm leaf, panicle flower, eating quality and tolerance are also significantly clustered together. Moreover, we performed the permutation test, discovering found that co-expression link densities between genes of same agronomic traits were significantly larger than random control gene set (Supplementary Text, Table S2). These results indicated that our gene networks can be used to discover the gene related to important agronomic traits by co-expression links.

### Function discovering for lncRNA genes

Long non-coding RNAs (lncRNAs) have been shown to play important roles in the kingdoms of plants and animals (Ranzani et al., 2015; Zhang et al., 2014). Given that the reconstructed RNA-seq-based co-expression network can successfully associate genes with biological functions and phenotypes of interest, we next wish to discover the functions for uncharacterized lncRNA genes using network-based method. We downloaded the known lncRNAs of rice identified in previous studies (Zhang et al., 2014). We then combined these lncRNA genes with MSU7.0 reference genes to establish co-expression network based on the ensemble inference pipeline. The obtained network is composed of 24875 genes, containing 24014 protein-coding gene and 861 lncRNA genes connected by 1357039 edges. Compared with the previous protein-coding gene network, 7692 novel protein-coding genes were captured and linked with 817 lncRNA genes. As there is no gold standard available to evaluate the predictive performance, we adopted gene-guide subnetwork analysis to illustrate the potential capacity of this network for lncRNA function discovering. We selected a well-studied lncRNA gene of *XLOC_057324,* which was verified involving in panicle development and fertility, to establish a gene subnetwork consisting of the two-step co-expression neighborhoods (Fig.5 and Dataset 7). In this subnetwork, 4 genes including *SSD1, PLA1, DEP1* and GSD1 related to panicle development or fertility. In addition, we also found that 7 genes (pink nodes) whose *Arabidopsis* homologs participate in meiosis, embryo development or reproductive process. According to the functional annotation, some genes (blue nodes) might be also involved in pollen development, such as two cyclin genes *CYCA2* and *CYCD2.* Interestingly, 3 lncRNAs of *XLOC_061753, XLOC_006119* and *XLOC_031878* expressed in the reproductive organs are contained in this subnetwork. These results are in good agreement with the experimentally verified role of *XLOC_057324.*

**Fig 5.**
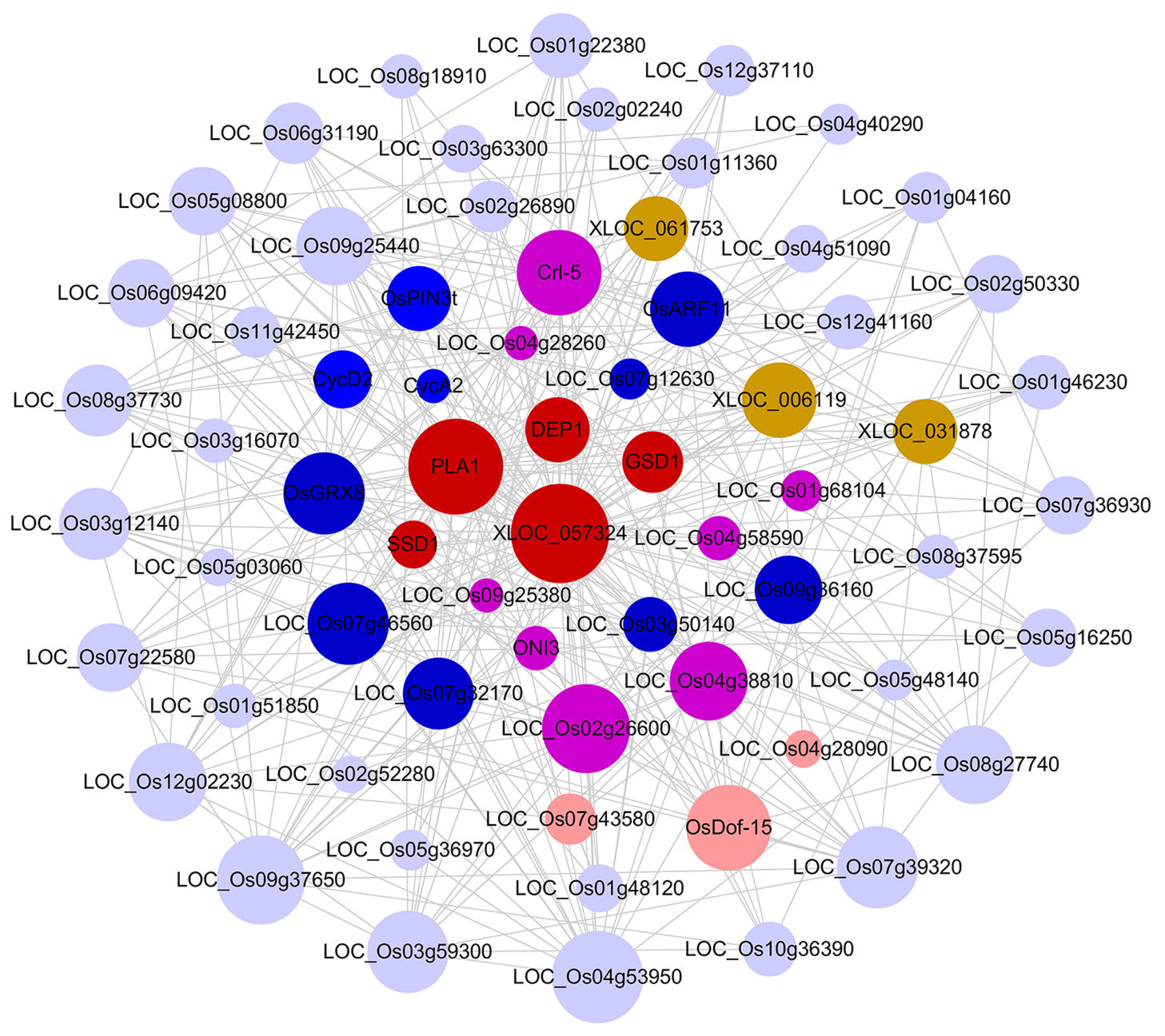
Co-expression subnetwork derived from guide-gene approach for *XLOC_057324* associated with panicle development and fertility. Within the subnetwork, red nodes represent the experimentally verifiedgenes related to corresponding biological functions; chrysoidine nodes represent transcription factors; pink nodes indicate the genes whose *Arabidopsis thaliana* homologues are experimentally verified to related to corresponding biological functions; blue nodes represent that the genes are potential function-related, and the gray nodes indicate that the genes are function unknown or annotated with unrelated functions

### CircRNA gene identification and function analysis

CircRNA is an RNA molecule forming a covalently closed continuous loop that has been discovered in various species across the domains of life with distinct sizes (Memczak et al., 2013; Ye et al., 2015). The functions of circRNAs are largely unknown and hard to investigate. Therefore, we try to classify them through gene co-expression network. We first identified 14325 circRNAs in rice derived from 5284 genes including 4609 protein-coding genes, 675 noncoding genes (see Materials and Methods for details). 43 of these genes including 27 protein-coding genes and 16 non-coding genes produce the circRNAs with the percentage larger than 90% in at least one sample. We analyzed the distribution of the number of detected circRNAs and found that a majority of circRNAs were identified in one sample with relative small number of circRNAs were detected in more than 3 samples (Fig.S14A). Though a large number of circRNAs were detected in relative small number of RNA-seq samples, 63 circRNAs (transcribed from the protein-coding genes), identified in more than 10 samples and supported by more than 26 junction reads, were captured in the gene co-expression network. Moreover, we found that the primary genes transcribing these circRNAs were not contained in the co-expression network. We predicted the functions of these circRNAs using GO enrichment analysis of their co-expression neighborhoods. Indeed, these circRNAs are related to a broad range of biological functions, for example protein phosphorylation, ATP binding and photosynthesis (Fig.S14B). These results indicated that a great number of circRNAs play important biological roles but not are the transcriptional noise.

## Discussion

The phenotypes of an organism are determined by the coordinated activity of many genes and gene products. To gain insight into the genetic foundation underlying the complex biological processes and phenotypes, we developed a novel analytic pipeline for constructing high-quality RNA-seq-based co-expression network and predicting gene function and regulations. we applied this pipeline to the important crop species rice. The obtained co-expression links between genes were ranked by confidence score, expression level and expression sample number. The thresholds of these measures can be selected as the indictors of co-expression reliability for the further targeted experimental validation. The detailed analysis of the topology properties of network demonstrates that the degree frequency distribution follows the truncated power-law and network structure is highly modular. Using the rice gold standards and bottom-up co-expression subnetwork analysis, we showed that this analysis pipeline can be effectively applied to study the gene function and regulation. Particularly, the potential application value of RNA-seq gene network for predicting biological roles of lncRNA and circRNA genes are well demonstrated. Overall, our analysis provides new insights into the regulatory code underlying transcription control and a starting point for understanding the complex regulatory system.

Compared with the sequence-based functional annotation, a great advantage of gene co-expression-based inference approach is that homologs are not required for a gene to receive a prediction. Actually, it is the case when a novel function appears for a particular species and the genes participating in the new biological process do not have corresponding homologues in other species. This is especially interesting for the non-coding RNAs because only short regions of non-coding RNA transcripts are limited by sequence- or structure-specific interactions, compared to the protein-coding gene; this difference in selection pressure makes it very difficult to find orthologous non-coding RNAs by their sequences. Indeed, using the BLAST search against NCBI Reference Sequence Database (RSD), we found that 87% and 89% of unannotated genes and lncRNA genes do not have homologous genes in other species, respectively. The functional analysis of rice lncRNA gene of *XLOC_057324* suggested that our RNA-seq-based gene network can be effectively applied to annotate the functions of non-codinggenome elements.

For RNA-seq-based gene co-expression network investigators, the creation of novel computational methods for building high-quality network poses a future fundamental challenge. According to our best knowledge, only four existing methods including Pearson’s Correlation Coefficient (PCCs), WGCNA, Canonical Correlation Analysis (CCA) and SpliceNet have been used to establish the RNA-seq gene co-expression networks (Giorgi et al., 2013; Hong et al., 2013; Iancu et al., 2012; Yalamanchili et al., 2014). Moreover, some of these inference tools are unable to be applied to the large-scale expression dataset owing to their high computational complexity. For the uncertainty and complexity of mechanism models underlying the RNA-seq data, we designed a novel ensemble-based inference pipeline to establish the high confidence RNA-seq gene co-expression network. Our outcomes demonstrate that the committee of three inference methods provides more robust and less false positive and false negative results than single algorithm. The improved performance of our ensemble inference method depends on the voting and rescoring scheme which can reduce the bias occurring in a single learning method and assign a higher confidence to the interactions that are repeatedly retrieved by different methods. Indeed, the standpoint of aggregating the results of different algorithms has been adopted in various contexts and it has proven to be effective in a variety of applications (Lertampaiporn etal., 2013; Liu et al., 2007; Yang et al., 2010).

In principle, gene co-expression meta-analysis can only detect co-regulations between genes which are co-expressed constantly or are sometimes co-expressed but otherwise silent. However, many activation patterns of gene groups appear only under the specific experimental conditions but behave independently under the other conditions, which might not be captured by our method. Especially, for lncRNA and circRNA genes, their expression patterns demonstrated highly spatiotemporal specificity. To overcome this problem, the high-efficiency bi-clustering methods can be integrated into our model to reveal the transcriptional gene interactions presented only under a specific subset of the experimental conditions (Madeira and Oliveira, 2004). Our approach can further improved by I) expanding our ensemble pipeline with other high-efficiency inference methods (Hase et al., 2013), II) employing more reasonable voting and rescoring schemes to generate the consensus networks.

## Materials and methods

### Dataset preprocessing

We downloaded 456 rice primary RNA-seq samples from the NCBI Sequence Read Archive (see Dataset 1 and 2 for details), with the keywords of “*Oryza sativa*” [Organism] AND “platform illumina” [Properties] AND “strategy rna seq” [Properties] (accessed on May 29, 2014). These RNA-seq samples contained a wide spread of experimental conditions, tissue types and developmental stages. After the SRA files were gathered, the archives were extracted and saved in FASTQ format using the SRA Toolkit. The FASTQ files were firstly trimmed using Trimmomatic software (version 0.32) (Bolger et al., 2014) with the default settings, except for an additional parameter of minimum read length at least 70% of original size. Then, the fastq_quality_filter program included in FASTX Toolkit was adopted to further filtrate the FASTQ files, with the minimum quality score 10 and minimum percent of 50% bases that have a quality score larger than this cutoff value. Surviving RNA-seq samples were mapped to the MSU7.0 reference genomes (55986 genes) using TopHat v2.0.4 with the default settings except for “‐‐max-multihits 1” (Trapnell et al., 2009). The PCR and optical/sequencing-driven duplicate reads were removed using the Picard tools. After reads mapping, the uniquely aligned reads count (RAW) and Fragments Per Kilobase Of Exon Per Million Fragments Mapped (FPKM) of each gene was calculated relative to the reference gene model using the HTSeq-count (v0.5.4) and Cufflinks software (v2.1.1), respectively (Anders et al., 2014; Trapnell et al., 2012). The unreliable samples and genes were filtered according to the following three criteria: I) The samples,in which the percentage of the number of genes with expression value smaller than 10 reads is larger than 90%, were not considered for further analysis; II) We did not consider the genes whose expression value is less than 10 reads in more than 80% samples; III) Genes with the variation coefficient of expression values smaller than 0.5 were excluded from subsequent analysis. After filtering, we got two expression datasets composed of 348 RNA-seq samples and 24775 genes were. The filtered RAW dataset were further corrected using four normalization methods: I) Upper Quartile (UQ) (Robinson et al., 2010); II) Trimmed Mean of M values (TMM) (Robinson et al., 2010); III) Relative Log Expression (RLE) (Robinson et al., 2010) and IV) Variance Stabilizing Transformation (VST) (Anders and Huber, 2010).

The microarray gene expression data were extracted from both ATTED-II database and Rice Oligonucleotide Array Database (ROAD) (Cao et al., 2012; Obayashi et al.,2009). The Gene Ontologies (GOs) were downloaded from Plant GeneSet Enrichment Analysis Toolkit(PlantGSEA) (Yi et al., 2013). We downloaded biological pathways from two data sources including PlantGSEA database and Plant Metabolic Network (PMN) (http://pmn.plantcyc.org/). The gene sets of transcription factor family were downloaded from Plant Transcription Factor Database (PlantTFDB) (Jin et al., 2013). MicroRNAs and their related targets were collected from the Plant MicroRNA Target Expressiondatabas(PMTED) and Plant MicroRNA database (PMRD) (Zhang et al.,2010).Known agronomic trait genes were collected from both Q-TARO database (Yonemaru et al., 2010) and literatures. Tos17 mutant phenotypes were extracted from Rice Tos17 Insertion Mutant Database (Hirochika et al., 1996). The phenotypes were associated with MSU7.0 gene locus identifiers through BLASTN alignments of Tos17 flanking sequences obtained from NCBI website. Protein-protein interaction network of rice were downloaded from PRIN (Gu et al., 2011). Probabilistic functional gene network of rice was obtained from RiceNet data portal (Lee et al., 2011).

### Gene co-expression network construction

We developed an ensemble-based inference pipeline for constructing the high-quality RNA-seq-based Gene Co-expression Network (GCN) based upon combining multiple inference algorithms, then aggregating their predictions through an unweighted voting system and rescoring co-expression links. Our ensemble-based inference system was designed based on the hypothesis that the different network inference methods have complementary advantages and limitations under the different contexts. To select base inference methods for constructing an ensemble system, five algorithms were initially tested and evaluated, including the weighted gene co-expression network analysis (Langfelder and Horvath, 2008), graphical Gaussian model (Schäfer et al., 2001), bagging statistical network inference (de Matos Simoes and Emmert-Streib, 2012), graphical lasso model (Friedman et al., 2008) and tree-based method (Huynh Thu et al., 2010). Since graphical lasso and tree-based method have high computational complexity and are infeasible for large number of RNA-seq dataset, we did not adopt these two algorithms for subsequent network construction. The flowchart for building high confidence RNA-seq-based gene co-expression network was depicted in Fig.6. In details, our procedure for producing the high-quality gene co-expression network was started from 6 RNA-seq datasets as described in Dataset preprocessing. Based on the 6 RNA-seq expression datasets, the weighted co-expression network inference, graphical Gaussian model and bagging statistical network inference were adopted to obtain 18 initial gene co-expression networks using the R packages of WGCNA, GeneNet and BC3NET, respectively (available from the CRAN repository). Since the outputs of WGCNA and GeneNet produced the long ordered list of confidence scores (topological overlap for WGCNA and partial correlation coefficient for GeneNet) for an enormous amount of gene pairs, we designed a random permutation model to choose the restrict threshold that roughly identifies functional co-expression links. We repeatedly created 100 times random datasets to obtain a series of background distributions, by randomly shuffling the associations from genes to expression profiles, and used the average of 99.99th percentile of these distributions (corresponding to the probability of 10^−4^ that two genes are connected by chance) to define the threshold. After obtaining initial networks, we employed two-step voting procedure, including voting within inference method and voting among the inference methods, to construct the high-quality gene co-expression network. In the first step of voting procedure, we selected the links included in more than two networks of all 6 initial co-expression networks, which were built by applying the single network inference algorithm to 6 RNA-seq datasets, to establish a consensus gene network (i.e. intra-method consensus network). In second step of voting procedure, we pick the co-expression relationships contained in more than one network of three intra-method consensus networks to establish the final co-expression network.

**Fig 6.**
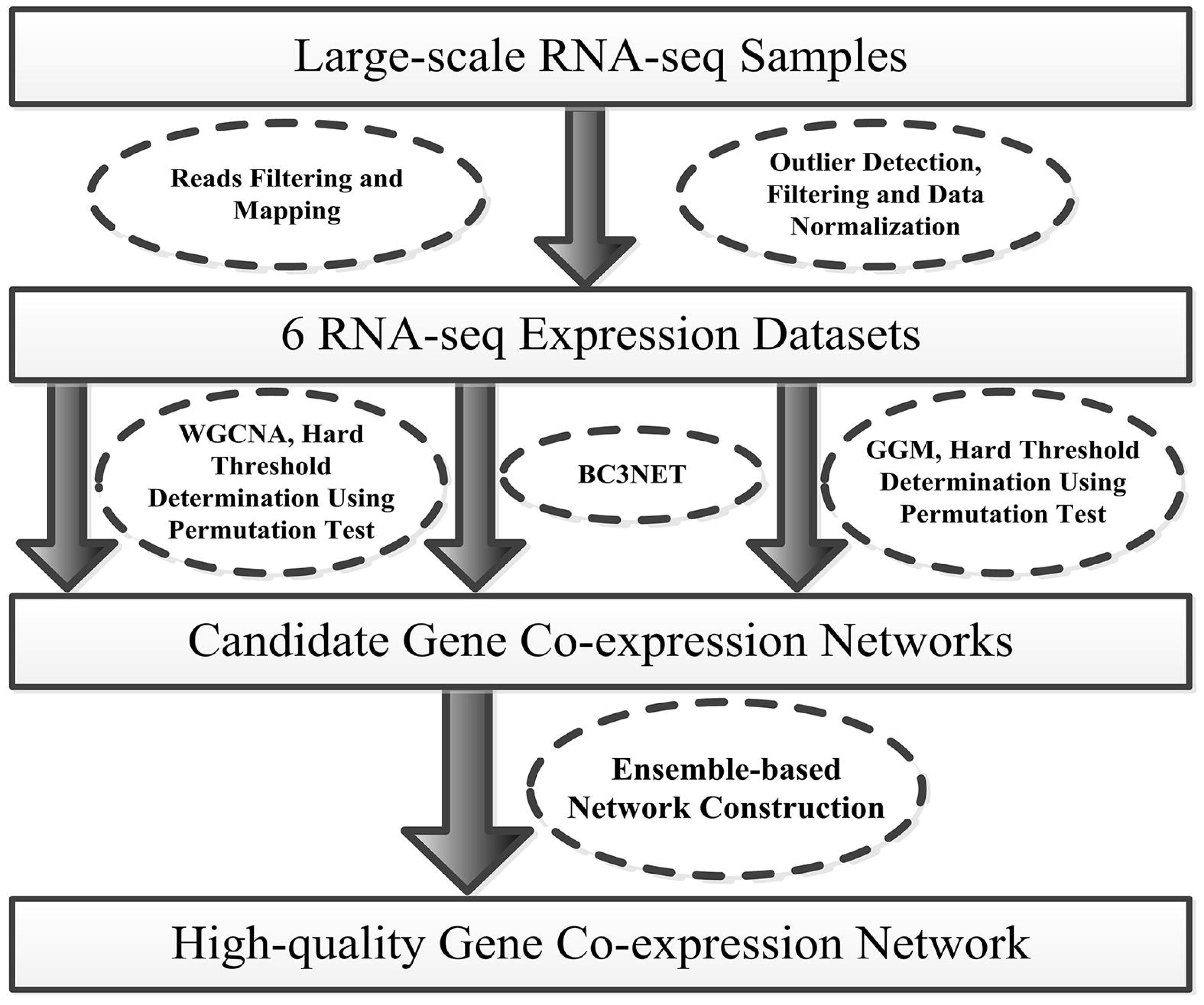
Flowchart of high-quality RNA-seq-based gene co-expression network inference

The confidence score calculation procedure for each gene pair of the final RNA-seq gene co-expression network was performed as following: I) Firstly, we normalized the confidence scores of each co-expression link of each initial network to the interval range from 0 to 1. II) Then, we assigned a confidence score to each association of the intra-method consensus gene networks by averaging the normalized confidence scores of all 6 initial networks. III) Finally, we defined the confidence score for each edge of final high confidence co-expression network by averaging the confidence scores of three intra-method consensus gene networks. Note that for the co-expression links not listed in a co-expression network were assigned a confidence score of 0.

### Performance evaluation

As the information about gold standard *Oryza sativa* reference gene network is unavailable, we compiled as replacement a standard set of positive and negative links for the performance evaluation. The gold standard of positive functional links was obtained by capturing gene pairs that were contained in the same GO categories, the same pathways, interact with each other in protein-protein interaction network or linked in probabilistic functional gene network. To construct the gold standard of negative functional links, we firstly selected all the biologically unrelated GO pairs (semantic similarity score = 0) that have the number of genes greater than 5 and less than 50, coupling all possible gene pairs of each partnership in remainder GO terms as initial non-functional relationships. Subsequently, we established 10000 background distributions of functional similarity, by 10000 times randomly sampling of 1000 gene pairs and calculating the functional similarities. We selected a subset of gene pairs from the initial non-functional links as final non-functional links using the criterion that the functional similarity between gene pair that are smaller than the average of 5th percentiles of these simulated background distributions. The semantic similarities between the GO terms were calculated using the R package of GOSim (Fröhlich et al., 2007). Functional similarities between genes in terms of the GO space were calculated using the metric adopted from (Chabalier et al., 2007).

Since our gold standards included only a subset of true functional and non-functional link, we evaluated the predictive performance of our method for gene co-expression network inference using the fold enrichment measure. The fold of enrichment was calculated as a function of the confidence score cutoff (*k*) in the edge list of the inferred network by the following formula:

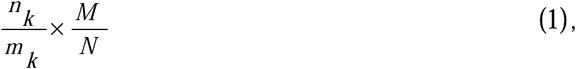

where, *n_k_* is the number of true positive or true negative functional links in the *k*th cutoff of the edge list; *m_k_* is the number of edges of the inferred network in the *k*th cutoff; *M* denotes the number of true positive or true negative functional links in the gold standards and *N* represents the number of all possible interactions in the genome space. The network visualization was carried out using both Cytoscape (Cline et al., 2007) and BioLayout Express3D (Theocharidis et al., 2009).

The function enrichment of co-expression neighborhoods was calculated as the ratio of the relative occurrence in gene set of co-expression neighborhoods to the relative occurrence in genome using Fisher’s exact test. The *p*-value was further adjusted by Benjamini-Hochberg correction for multiple hypotheses testing. The corrected *p*-value smaller than 0.05, was considered as enriched. To evaluate the predictive performance of our RNA-seq-based network for inferring gene function using the co-expression neighborhoods, we adopted the gene-centric evaluation, which were provided in the Critical Assessment of protein Function Annotation (CAFA) project (Tzafrir et al., 2003). For this metric, the GO terms of each gene (gold and predicted) are propagated up the GO hierarchy to the root, obtaining a set of terms. In this process, for each scored GO term, we propagated its score (-log(ç-value) of Fisher’s exact test) toward the root of the ontology such that each parent term received the highest score among its children. The Sensitivity (Recall), 1-specificity, Precision and maximum F-measure (F-max) was calculated using the same method as in the CAFA project. The Receiver Operating Characteristics (ROC) curve was drawn by changing the threshold and plotting the Sensitivity versus the 1-specificity and then calculated the score of Area Under Curve. Similarly, we plotted the Precision-Recall (PR) curve by altering the threshold and plotting the Precision versus the Recall. Semantic similarity scores between the GO term pairs were calculated using the R package of GOSim.

### Analysis of circRNA genes

The circular RNA (circRNA) genes were predicted using 618 novel rice RNA-seq samples downloaded from the NCBI Sequence Read Archive (accessed on February 15, 2016) by CIRI software (Gao et al., 2015). We calculated the counts of junction reads of a circRNA as its relative expression abundance. Then, we integrated the aligned reads number of known rice genes using HTSeq-count program (v0.5.4) and expression values of circRNAs into a numeric expression matrix. We removed the circRNAs from the matrix if it was identified in less than 3 RNA-seq samples. Using the filtered matrix, we built three initial gene co-expression networks by WGCNA, GGM and BC3NET. Based on this, we selected the co-expression links contained in more than one network of the three initial networks to obtain the final co-expression network. Although only the numbers of junction reads were adopted to measure the expression abundances of circRNAs, this method is simple and effective for building co-expression network, given the reads were distributed uniformly along circRNA.

## Acknowledgements

This work is financially supported by The Strategic Priority Research Program of the Chinese Academy of Sciences (Grant No. XDA08020302). The funders had no role in study design, data collection and analysis, decision to publish, or preparation of the manuscript.

## Author Contributions

H.Y. conceived the original screening and research plans; H.Y. and C.Z.L conceived the project; C.Z.L. and H.Y. supervised the experiments; H.Y. performed most of the experiments, analyzed the data and wrote the paper; B.K.J analyzed the phenotype data and revised the paper.

## Additional Information

**Competing financial interests:** The authors declare no competing financial interests.

